# Hybrid ssDNA repair templates enable high yield genome engineering in primary cells for disease modeling and cell therapy manufacturing

**DOI:** 10.1101/2021.09.02.458799

**Authors:** Brian R. Shy, Vivasvan Vykunta, Alvin Ha, Theodore L. Roth, Alexis Talbot, David N. Nguyen, Yan Yi Chen, Franziska Blaeschke, Shane Vedova, Murad R. Mamedov, Jing-Yi Chung, Hong Li, Jeffrey Wolf, Thomas G. Martin, Lumeng Ye, Justin Eyquem, Jonathan H. Esensten, Alexander Marson

## Abstract

CRISPR-Cas9 offers unprecedented opportunities to modify genome sequences in primary human cells to study disease variants and reprogram cell functions for next-generation cellular therapies. CRISPR has several potential advantages over widely used retroviral vectors including: 1) site-specific transgene insertion via homology directed repair (HDR), and 2) reductions in the cost and complexity of genome modification. Despite rapid progress with *ex vivo* CRISPR genome engineering, many novel research and clinical applications would be enabled by methods to further improve knock-in efficiency and the absolute yield of live knock-in cells, especially with large HDR templates (HDRT). We recently reported that Cas9 target sequences (CTS) could be introduced into double-stranded DNA (dsDNA) HDRTs to improve knock-in, but yields and efficiencies were limited by toxicity at high HDRT concentrations. Here we developed a novel system that takes advantage of lower toxicity with single-stranded DNA (ssDNA). We designed hybrid ssDNA HDRTs that incorporate CTS sites and were able to boost knock-in percentages by >5-fold and live cell yields by >7-fold relative to dsDNA HDRTs with CTS. Knock-in efficiency and yield with ssCTS HDRTs were increased further with small molecule inhibitor combinations to improve HDR. We demonstrate application of these methods across a variety of target loci, knock-in constructs, and primary human cell types to reach ultra-high HDR efficiencies (>80-90%) which we use for pathogenic gene variant modeling and universal gene replacement strategies for *IL2RA* and *CTLA4* mutations associated with mendelian immune disorders. Finally, we develop a GMP-compatible method for fully non-viral CAR-T cell manufacturing, demonstrating knock-in efficiencies of 46-62% and generating yields of >1.5 x 10^9^ CAR+ T cells, well above current doses for adoptive cellular therapies. Taken together, we present a comprehensive non-viral approach to model disease associated mutations and re-write targeted genome sequences to program immune cell therapies at a scale compatible with future clinical application.

## Introduction

CRISPR-Cas9 genome edited human cellular therapies have recently entered the clinic. Cas9-based knock-outs in T cells and hematopoietic stem cells (HSCs) have demonstrated a promising safety profile and, in some cases, profound efficacy^1,2^. Forthcoming trials are now poised to introduce Cas9-mediated knock-ins by homology-directed repair (HDR) for correction of pathogenic mutations or insertion of novel therapeutic constructs^3–5^. In comparison to randomly integrating viruses or transposon-based approaches, Cas9-stimulated HDR allows for precisely targeted genomic changes which can improve the quality, uniformity, and safety of cellular products^6,7^. In addition to reducing potential integration risks, targeted genome editing can repurpose endogenous genetic circuits and eliminate the need for artificial promoters. This can have important functional benefits as demonstrated for targeted Chimeric Antigen Receptor (CAR) insertion into the *TRAC* locus (T cell Receptor Alpha chain Constant region), which enhances CAR-T cell potency and persistence in preclinical studies by taking advantage of the endogenous gene regulatory elements governing normal TCR expression^6^. The ability of HDR to mediate high efficiency and scarless insertion of these large multi-kilobase DNA constructs is unmatched currently by alternative precision genome editing tools such as base editors, prime editors, transposase, or recombinase approaches^8^. Introduction of large DNA sequence payloads will be essential for manufacturing many future clinical products including CAR-T cells and therapeutic gene replacement strategies, and provides the flexibility needed for the next generation of synthetic biology constructs^6,9,10^.

*Ex vivo* CRISPR genome editing of primary human T cells has been optimized extensively by our group and others, generally using electroporation of pre-complexed Cas9 and guide RNA (gRNA) ribonucleoproteins (RNPs) to generate targeted genomic breaks^6,11,12^. To introduce targeted sequence insertions or replacements with HDR, an HDR template (HDRT) is included that encodes the desired genetic change in between homology arms that flank the genomic break. Several different methods are used to introduce the HDRTs including viral transduction with recombinant adeno-associated virus (rAAV) or co-electroporation with naked DNA in double-stranded (dsDNA), single-stranded (ssDNA), circular, or linear formats^6,12,13^. Both the efficiency of HDR and the cellular toxicity correlate directly with the concentration and format of the HDRTs. For large constructs, rAAV-based methods have thus far achieved the most impressive knock-in efficiencies while maintaining minimal toxicity^14,15^. While rAAV vectors have led to rapid advances, incorporation for research and clinical use has been slowed by the cost and complexity of manufacturing these reagents. Co-electroporation of naked DNA has the potential to greatly increase the pace of innovation in gene modified cell therapies, since it can be done at a fraction of the cost and time required for viral vector development. Non-viral approaches have been applied within primary human cell types, however, further improvements are needed, especially for large templates, to reduce DNA toxicity, improve knock-in purity and cell yields, and advance towards clinical applications^11,12^.

We recently developed a method to enhance the knock-in efficiency of dsDNA HDRTs through incorporation of Cas9 Target Sequences (CTS), allowing the coelectroporated RNPs to bind the HDRTs and facilitate their delivery^11^. We found that knock-in efficiencies were substantially increased but with concurrent increases in cellular toxicity. This toxicity could be attenuated, but not eliminated, by inclusion of anionic polymers such as polyglutamic acid (PGA) to improve cell yields. In comparison to dsDNA, ssDNA exhibits less toxicity^12^. Cas9 binds to dsDNA targets, so we set out to establish an approach to adapt CTS-based enhancement of HDR to ssDNA templates. Here we have developed a hybrid HDRT using a long ssDNA with short regions of dsDNA containing CTS sites on each end. For simplicity, we refer to these hybrid HDRTs as ssCTS templates and refer to the fully double-stranded variants as dsCTS templates.

We discovered that ssCTS templates significantly increased knock-in efficiency while minimizing toxicity across a range of construct sizes, genomic loci, and clinically relevant cell types including primary human T cell subsets, B cells, NK cells, and HSCs. In addition, we evaluate a panel of small molecule inhibitors reported to enhance HDR in primary human T cells, identifying the optimal combinations and concentrations that work to further enhance HDR with ssCTS templates. Combining ssCTS templates with small molecule inhibitors, we achieve extremely high knock-in efficiencies that in some cases approach pure populations of HDR-edited cells (>80-90%) across a range of clinically relevant target sites, and we demonstrate the application of this approach for gene replacement strategies and functional evaluation of patient mutations. Finally, we adapt our approach to generate a GMP-compatible process for fully non-viral CAR-T cell manufacturing, achieving knock-in efficiencies of 40-62% at clinical-scale with production of >1.5 x 10^9^ CAR+ cells from a starting population of 100 x 10^6^ T cells. This technology should broadly enable wide-spread efforts to model patient mutations in primary cells and flexibly engineer cellular therapies at clinical-scale.

## Results

### Development of ssCTS templates for high-efficiency and low-toxicity HDR in primary human T cells

We previously developed a method to enhance delivery of dsDNA HDRTs through incorporation of Cas9 target sites (CTS) which include a gRNA target sequence and an NGG Protospacer-Adjacent-Motif (PAM) on each end of the template^11^. In comparison to dsDNA, ssDNA is associated with lower toxicity, which we reasoned could further improve knock-in efficiency and cell yield with large DNA templates if combined with CTS technology^12^. We screened a variety of hybrid structures composed predominantly of ssDNA with small stretches of dsDNA incorporating the CTS sites through hairpin loops, annealed complementary oligonucleotides, or more complex secondary structures (Fig. 1A-C)^12^. We rapidly screened to compare HDRT designs using short 113–195nt HDRTs that generate an N-terminal CD5-HA fusion protein easily detectable by flow cytometry (Supplementary Fig. 1A-B). We found that the majority of these ssCTS designs increased knock-in efficiency (Fig. 1C). Improved efficiency with the ssCTS templates was apparent only at the lower 2 concentrations (160nM and 800nM), above which the knock-in efficiency appeared to hit a maximum of ~30% that was achievable with unmodified ssDNA HDRTs (Fig. 1C, grey). These results suggested that ssCTS designs would be beneficial in situations where the HDRT concentration is limited, such as with large HDRTs that typically reach toxicity in the 10-320 nM range depending on their length and format.

**Figure 1.**
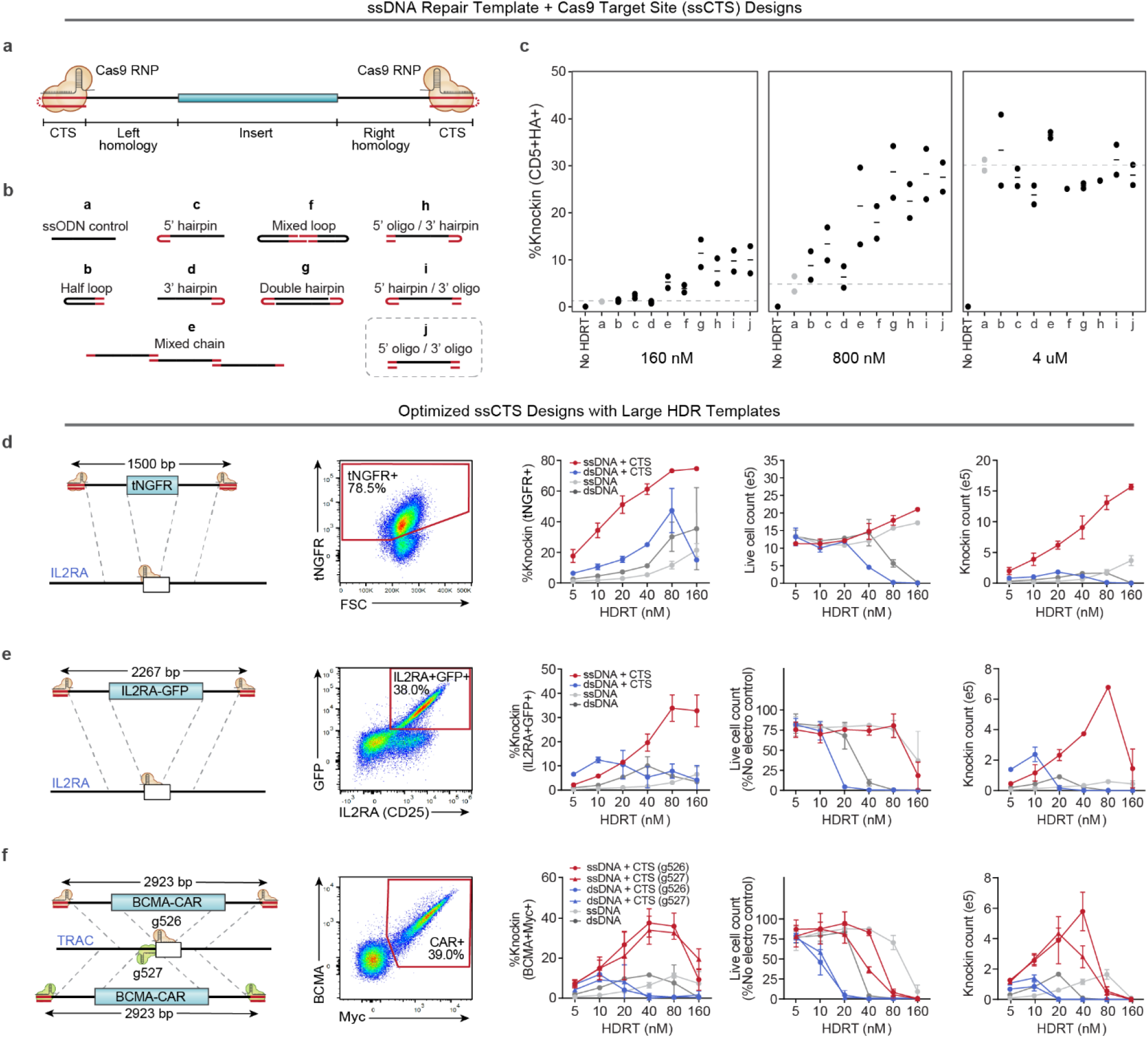
Development of ssCTS templates for high yield knock-in. **(a)** Diagram of hybrid ssDNA HDRT designs incorporating CTS sites (ssCTS). **(b)** Panel of ssCTS designs tested. **(c)** Knock-in efficiency for each ssCTS design using a CD5-HA knock-in construct at 160 nM - 4uM concentration assessed by flow cytometry. Dotted line represents mean knock-in percentage for control ssDNA HDRTs without CTS (construct a, grey). **(d-f)** Knock-in strategy, gating, knock-in efficiency, live cell counts, and knock-in cell counts are shown for large ssCTS templates including **(d)** a tNGFR knock-in at the *IL2RA* locus, **(e)** a IL2RA-GFP fusion protein knock-in to the *IL2RA* locus, or **(f)** two different HDRTs inserting a BCMA-CAR construct at *TRAC* locus via two different gRNAs (g526 and g527). Each experiment was performed with T cells from 2 independent healthy human blood donors. Error bars indicate standard deviation. RNP = Ribonucleoprotein, CTS = Cas9 Target Site, ssCTS = ssDNA HDRT + CTS sites, HDRT = homology-directed-repair template.

For evaluation of large HDRTs, we chose an ssCTS design that incorporates CTS sites on both the 5’ and 3’ end via annealed complementary oligonucleotides, which are easy to design for research and clinical applications. In our panel of tested ssCTS constructs, this design demonstrated maximal enhancement of knock-in efficiency (Fig. 1B-C, “j”), low toxicity (Supplementary Fig. 1C), and provided the simplest process for generating CTS ends compared to hairpin loops or more complicated structures. Long ssDNA and dsDNA HDRTs ranging from 1500nt to 2923nt were generated with and without CTS sites (Fig. 1D-F). These templates target a knock-in detectable by flow cytometry (tNGFR, IL2RA-GFP fusion, or BCMA-CAR) to the *IL2RA* or *TRAC* locus. We evaluated post-electroporation knock-in efficiency, toxicity (based on live cell counts), and absolute yield of successful knock-in counts using primary T cells isolated from healthy human blood donors. Inclusion of CTS sites enhanced the knock-in efficiency of both dsDNA and ssDNA constructs across concentrations until toxic doses were reached, after which knock-in efficiency progressively decreased. ssCTS constructs demonstrated uniformly higher knock-in efficiencies and lower toxicity in comparison with dsCTS templates, generating up to 7-fold more knock-in cells at optimal non-toxic concentrations. The use of ssCTS templates allowed us to achieve up to 78.5% knock-in with a ~1.5kb tNGFR construct, or 38% for a ~2.3kb IL2RA-GFP construct targeting the *IL2RA* locus; and up to 39% knock-in with a ~2.9kb BCMA-specific CAR construct targeting the TRAC locus at HDRT concentrations compatible with high yields of live knock-in cells.

### Exploration and optimization of ssCTS design parameters for large HDRTs

To learn rules regarding the precise sequences required for ssCTS-enhanced HDR, we evaluated variations of two constructs targeting either an IL2RA-GFP fusion to the *IL2RA* gene (~2.3kb, Fig. 1E) or a large version of the CD5-HA knock-in including >1kb homology arms (~2.7kb, Supplementary Fig. 2A). We first evaluated the specificity of the CTS sequences by replacing them with a mismatched CTS site specific for the alternative RNP, an equivalent length of dsDNA within the homology arm (“end protection”), or a CTS site with scrambled gRNA sequence (Fig. 2A, Supplementary Fig. 2B). For both constructs, only the matching CTS recognized by the cognate RNP increased knock-in efficiency, suggesting specific recognition of the gRNA sequence.

**Figure 2.**
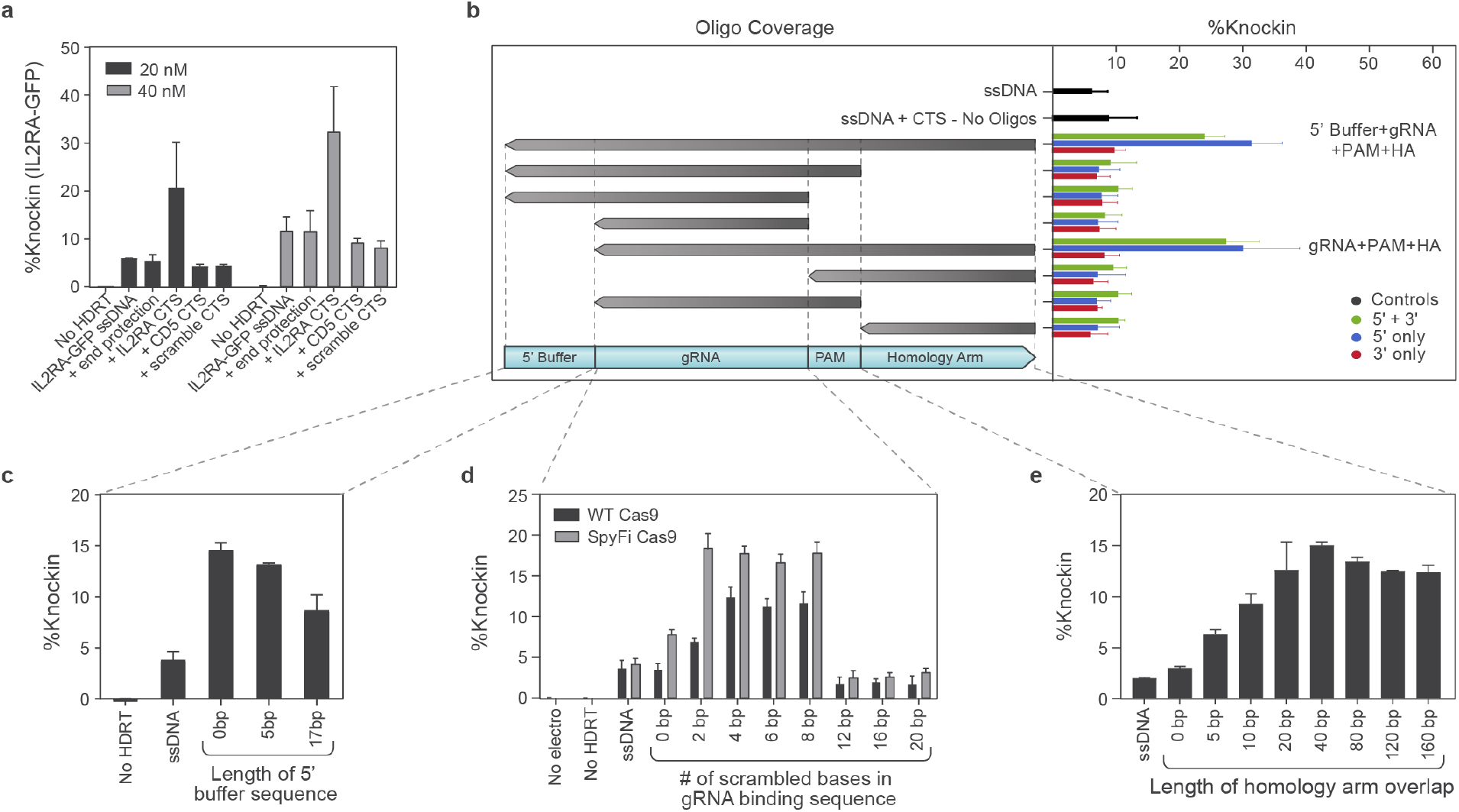
Evaluation of ssCTS design requirements. **(a-e)** Comparison of different CTS designs with an IL2RA-GFP knock-in construct targeting *IL2RA* locus assessed by flow cytometry. **(a)** Comparison of CTS with a gRNA target sequence that is specific for the cognate RNP (+ IL2RA CTS), an alternative gRNA sequence (+ CD5 CTS), a CTS incorporating a PAM site and scrambled gRNA sequence (+ scramble CTS), or an equivalent amount of dsDNA within the 5’ end of the homology arm (+ end protection). **(b)** Comparison of complementary oligos covering varying regions of the CTS and surrounding sequences (design schematics left; knock-in results right). Constructs with CTS sites on both 5’ and 3’ end (green bars), 5’ end only (blue blurs), or 3’ end only (red bars) are shown on the right panel with two best performing designs indicated (right). **(c)** Evaluation of varied 5’ ends including different length of buffer sequence upstream of the CTS site. **(d)** Comparison of CTS designs with varying numbers of scrambled bases at the 5’ end of the gRNA target sequence using WT or SpyFi Cas9. **(e)** Knock-in percentages are shown with varying length of homology arm covered by the complementary oligonucleotide. Each experiment was performed with T cells from 2 independent healthy donors. Error bars indicate standard deviation. All comparisons except for panel b include complementary oligos covering the entire 5’ Buffer + gRNA + PAM + homology arms. CTS = Cas9 Target Site, PAM = Protospacer Adjacent Motif, HDRT = homology-directed-repair template, HA = homology arms.

We next examined closely which components of the CTS required dsDNA by annealing oligos of varied lengths and coverage (Fig. 2B, Supplementary Fig. 2C). Coverage of the gRNA sequence, PAM, and a stretch of nucleotides within the homology arm downstream of the CTS site were each required for enhancement of knock-in efficiency while coverage of nucleotides upstream of the gRNA sequence in the 5’ buffer region was not. In agreement, inclusion of additional buffer sequence upstream of the CTS was not required at all and may in fact reduce the knock-in efficiency (Fig. 2C, Supplementary Fig. 2D). Surprisingly, we saw that inclusion of a CTS on the 3’ end of both large ssCTS constructs provided no independent or additive benefit in combination with a 5’ CTS. Similar findings were seen within our short HDRT screen (Fig. 1B-C, “c” versus “d”). These intriguing results suggest only the 5’ CTS is functional in these designs which could reflect requirements for RNP binding and orientation, intracellular trafficking, or interference with repair machinery during 3’ annealing of long ssDNA^16^.

We further examined the requirements for gRNA recognition by generating CTS sites with a variable number of scrambled bases at the 5’ end of the 20bp gRNA recognition sequence (Fig. 2D, Supplementary Fig. 2E). We found that for WT Cas9, the enhancement in knock-in efficiency was maximal with inclusion of 4-8 mismatched nucleotides. This level of mismatch likely allows the Cas9 RNP to bind without cleaving the CTS, as has been shown for truncated gRNAs^17^. The pattern was similar with the high fidelity “SpyFi” Cas9 variant produced by Aldevron/IDT, which has been developed to reduce off-target cuts in clinical gene editing applications^18^. Finally, we evaluated the length of the complementary oligonucleotide coverage within the downstream homology arm, demonstrating optimal knock-in when > 20-40bp of the homology arm has complementary sequence in the corresponding oligo (Fig. 2E, Supplementary Fig 2F). Taken together, these data establish design rules for end oligos to introduce CTS into large ssDNA templates and boost HDR outcomes, demonstrating that optimal designs need only incorporate a single CTS site on the 5’ end with a short stretch of dsDNA covering the gRNA recognition site, the PAM sequence, and ~20bp of the downstream homology arm (Fig. 2B, Supplementary Fig. 2C).

### ssCTS templates provide a flexible and powerful approach to enhance HDR in primary human cells

Using optimized ssCTS designs, we next assessed performance across a broad array of genomic loci, knock-in constructs, and primary hematopoietic cell types. We first evaluated an arrayed panel of knock-in constructs in primary human T cells targeting a detectable tNGFR fusion at the 5’ end of 22 different genes (Fig. 3A). The majority of ssCTS constructs outperformed alternative HDRT variations for both knock-in efficiency (up to 5-fold increase) and absolute knock-in counts (up to 3-fold increase) with only a few exceptions that appeared equivalent to dsCTS constructs (Fig. 3A, Supplementary Fig. 2G). We next evaluated performance with a pooled library of knock-in constructs targeting an NY-ESO-1 specific TCR and additional gene products to the endogenous *TRAC* locus, as previously reported by our group for use in functional knock-in screens^19^ (Fig. 3B-D). Knock-in pools provide a powerful approach for high-throughput screening and allowed us to assess performance with a diverse population of large knock-in templates ranging from 2.6-3.6kb. Knock-in efficiency and absolute knock-in counts were both increased by >5-fold in comparison to optimal dsCTS concentrations, significantly increasing coverage for each individual construct while retaining consistent representation of the initial library in the final knock-in population (Fig. 3B-D). Finally, we evaluated performance across a variety of clinically relevant primary cell types including CD4+ T cells, CD8+ T cells, regulatory T cells (Treg), NK cells, B cells, hematopoietic stem cells (HSC) and gamma-delta T cells using an mCherry knock-in construct targeting the clathrin light chain A (*CLTA*) gene (Fig. 3E-G). In all evaluated cell types ssCTS templates demonstrated significantly lower toxicity, increased knock-in efficiency, and generated higher absolute knock-in cell yields.

**Figure 3.**
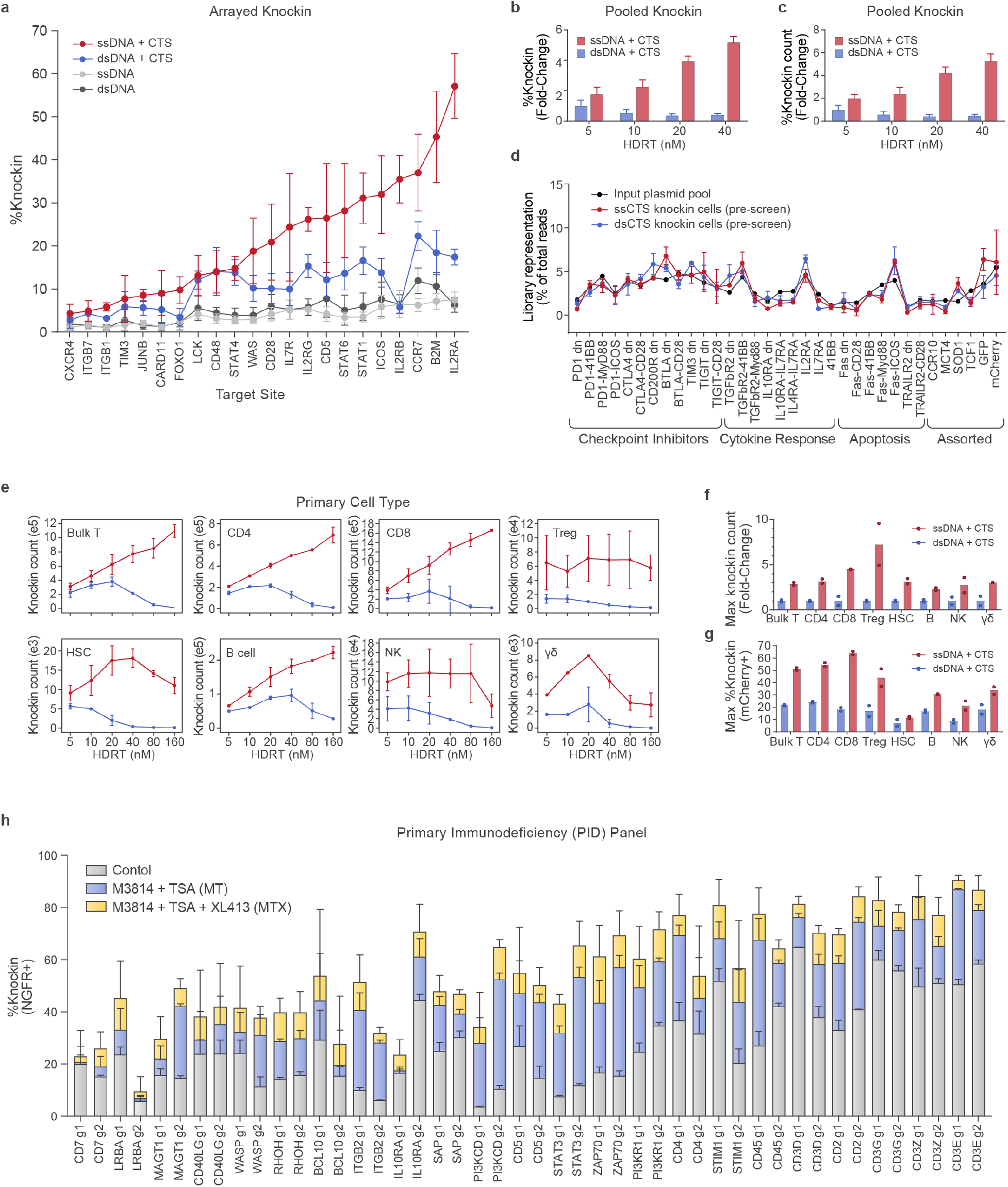
Application of ssCTS knock-in templates across diverse target loci, knock-in constructs, and primary human hematopoietic cell types. **(a)** Knock-in efficiencies for constructs targeting a tNGFR marker to 22 different target genome loci. **(b-d)** Comparison of large ssDNA and dsDNA HDRTs with CTS sites for knock-in of a pooled library of 2.6 - 3.6 kb polycistronic constructs targeted to the *TRAC* locus^19^. Shown for each HDRT variation is **(b)** relative %knock-in in comparison to maximum for dsDNA + CTS templates, **(c)** relative knock-in cell count yields in comparison to maximum for dsDNA + CTS templates, and **(d)** representation of each library member in knock-in cells post-electroporation in comparison to construct representation the input plasmid pool. **(e-g)** Comparison of knock-in cell yields using ssDNA (red) and dsDNA HDRTs (blue) with CTS sites across a variety of primary human hematopoietic cell types. Note, different cell type comparisons are performed with different blood donors. All comparisons were performed using a knock-in construct generating an CLTA-mCherry fusion at the *CLTA* locus. Shown for each cell type using HDRT concentrations from 5-160 nM are **(e)** knock-in cell count yields, **(f)** maximum fold-change in knock-in count yields (relative to dsCTS templates), and **(g)** maximum %knock-in. **(h)** Evaluation of ssCTS templates +/- M3814 + TSA (MT) or M3814 + TSA + XL413 (MTX) inhibitor combinations with a panel of 44 different knock-in constructs targeting a tNGFR marker across 22 different target loci including genes implicated in Primary Immunodeficiencies (PID) or with potential importance for T cell engineering. 2 gRNAs and corresponding ssCTS templates were used for each gene (g1 and g2). All experiments were performed with T cells from 2 independent healthy donors. Error bars indicate standard deviation. CTS = Cas9 Target Site, HDRT = homology-directed-repair template, tNGFR = truncated Nerve Growth Factor Receptor, dsCTS = dsDNA HDRT + CTS sites, ssCTS = ssDNA HDRT + CTS sites, kb = kilobase, MT = M3814 + TSA, MTX = M3814 + TSA + XL413.

### Evaluation of small molecule inhibitors in combination with ssCTS templates for high-efficiency T cell engineering at Primary Immunodeficiency (PID) disease loci

We next evaluated a panel of small molecule inhibitors reported to enhance knock-in efficiency in primary human T cells including the DNA-PK inhibitors NU7441 and M3814, the HDAC class I/II Inhibitor Trichostatin A (TSA), the CDC7 inhibitor XL413, and IDT’s proprietary Alt-R HDR Enhancer which is described as a NHEJ inhibitor^20–22^. Using our short ssDNA CD5-HA knock-in construct (Fig. S1A-B), each was titrated in isolation and then evaluated in combination to identify effects on knock-in efficiency and live cell counts (Supplementary Fig. S3A-B). At optimal concentrations, M3814 showed the largest effect size (~49% increase), followed by XL413 (~46% increase), NU7441 (~43% increase), IDT’s HDR Enhancer (~29% increase), and TSA (~16% increase). Live cell counts were generally unaffected at the chosen concentrations except for combinations involving XL413, which demonstrated an ~50% reduction in cell counts at day 4 post-electroporation that may reflect XL413’s mechanism as a transient cell cycle inhibitor rather than overt cytotoxicity (Supplementary Fig. 4B)^22^. NHEJ inhibitor combinations (M3814, NU7441, IDT HDR Enhancer) did not demonstrate further improvements above the highest individual component, consistent with overlapping mechanisms of action. In contrast, addition of TSA or XL413 did demonstrate additional improvements in combination with NHEJ inhibitors. The M3814/TSA (MT) combination provided the largest increase in knock-in efficiency without affecting live cell counts (~65% increase) and the M3814/TSA/XL413 (MTX) combination demonstrated the highest absolute increase in knock-in efficiency (~134% increase) albeit with XL413-mediated reduction in total cell counts. Finally, we evaluated whether the benefits of ssCTS templates and small molecule inhibitors could be combined using a variety of constructs ranging from 1.5-2.7kb (Supplementary Fig. 3C). Encouragingly, each construct demonstrated increased knock-in efficiency with ssCTS templates that was further enhanced by the inclusion of MT and MTX inhibitor combinations, in some cases generating knock-in efficiencies >90%.

Encouraged by these results, we sought to evaluate these approaches more broadly and at clinically relevant target sites that could lead toward diagnostic or therapeutic advances. We developed a panel of knock-in constructs targeting genes associated with monogenic disease-causing mutations affecting T cell function or relevant controls. These diseases are part of a spectrum of increasingly recognized genetic disorders, referred to as Primary Immunodeficiencies (PID) or Inborn Errors of Immunity (IEI), that disrupt the healthy immune system, presenting with severe infections, autoimmune disease, and malignancy^23^. Within this panel, we examined 44 different tNGFR constructs targeting 22 genes (2 gRNA targets per gene) using ssCTS templates +/- MT and MTX inhibitor combinations (Fig. 3H). This analysis demonstrated nearly universal increases in knock-in efficiency with MT that were further enhanced with the MTX combination, achieving knock-in rates >50% for these large constructs at 15/22 genes examined and >80% at 6/22 genes. The effect size of inhibitors varied among target loci, with some sites demonstrating relatively little increase (e.g. CD7 g1) and others showing up to 7-fold increases (e.g. PI3KCD g2). Live cell counts were comparable at day 5 post-electroporation with a few notable exceptions demonstrating significant toxicity with both combinations (e.g. CD7 g2, WASP g2, CD3G g2) (Supplementary Fig. 3D). Altogether these findings support broad application of ssCTS templates and inhibitor combinations at relevant disease loci, in some cases demonstrating nearly pure populations of knock-in cells (>80-90%) (Fig. 3H, Fig S3C). This sets the stage for diagnostic and therapeutic applications of non-viral human T cell engineering that require a high purity or yield of knock-in cells at specific disease loci.

### Universal gene replacement strategies for therapeutic and diagnostic human T cell editing

To explore potential clinical applications with large non-viral templates, we chose to examine whole open reading frame (ORF) insertions for two genes, *IL2RA* and *CTLA4*, that have been identified in families with monogenic immune disorders characterized by severe multi-organ autoimmunity^12,24–31^. Although disease-causing mutations are widely distributed throughout these genes, many of these families could potentially be treated by a universal ORF replacement strategy (Fig. 4A, 4E). For each construct, we included a GFP fusion at the 3’ end to facilitate detection of the knock-in protein. We have previously reported targeted gene corrections for a family with loss-of-function mutations in exon 4 and exon 8 of the *IL2RA* gene^12^. While we achieved knock-in efficiencies >30% with this approach, each site required a custom gRNA and HDRT which prevents extension to families with alternative *IL2RA* mutations. In contrast, a whole ORF knock-in at exon 1 of the *IL2RA* gene could potentially ameliorate any of the 11 previously reported mutations causing IL2RA deficiency (Fig. 4A)^12,24–28^. Using a ssCTS template and the MTX inhibitor combination, we achieved >80% knock-in of a ~2.3kb whole ORF IL2RA-GFP fusion construct (Fig. 4B). The knock-in protein demonstrated nearly indistinguishable expression levels compared to endogenous protein. This whole ORF knock-in approach could allow for rapid functional testing and characterization of patient mutations or variants of unknown significance (VUS). To demonstrate this diagnostic potential, we modified the knock-in construct to encode a previously described disease-causing mutation in exon 4 of *IL2RA*, c.497 G>A (S166N), which was reported to eliminate surface expression while retaining cytoplasmic protein^27^. In agreement with what has been reported in patient cells, we found that the GFP+ S166N population demonstrates a near complete absence of surface IL2RA with readily detectable intracellular IL2RA comparable to WT levels (Fig. 4C, Supplementary Fig. 4A). Fluorescence microscopy revealed that S166N protein formed distinct perinuclear aggregates consistent with intracellular retention and contrasting with the diffuse cytoplasmic and surface IL2RA seen with WT knock-ins (Fig. 4D). These results highlight the diagnostic and therapeutic potential of targeted ORF insertion within the endogenous gene, an approach which may be extended to include a number of alternative targets or additional noncoding elements.

As a further example, we examined an ORF insertion within the *CTLA4* gene (Fig. 4E). CTLA4 deficiency is caused most frequently by a haploinsufficiency with a disease-causing mutation on only 1 of 2 alleles ^29–31^. Exon-targeting strategies generate indels which could disrupt the normal allele and worsen disease. To avoid this possibility, we screened a panel of gRNA in intron 1 to identify targets which cut efficiently without disrupting protein expression (Supplementary Fig. 4B). The chosen gRNA had no detectable disruption of endogenous CTLA4 protein and the associated ORF knock-in construct generated knock-in efficiencies of 70-80% with ssCTS templates and MTX inhibitor combination (Fig. 5F-G). This intron-targeting strategy could be used to introduce or correct the majority of reported disease-causing mutations in *CTLA4* excluding those upstream of the target site (Fig. 5E). Variations in protein expression by cell type and in response to stimulation matched the endogenous protein, although basal knock-in protein levels were slightly higher which may reflect differences between the SV40 3’UTR used in this construct and the endogenous 3’UTR (Supplementary Fig. 4C)^32^. To evaluate diagnostic capabilities with known *CTLA4* mutations, we generated knock-in constructs with 3 previously reported disease-causing mutations: R70W, R75W, and T124P^29^. Cells were gated for the highest levels of GFP expression to enrich for homozygous knock-ins and then evaluated for surface protein, intracellular protein, and ligand binding using recombinant CD80 in activated CD4+ T cells (Supplementary Fig. 4D, Fig. 4G-I). All three mutations significantly reduced ligand binding despite variable levels of surface expression, in agreement with prior reports demonstrating reduced ligand uptake in heterozygous patient cells or engineered cell lines^29^. Altogether, these approaches provide a powerful method for evaluating patient mutations at endogenous loci with the potential for adaptation to high-throughput screening and high efficiency therapeutic gene replacement strategies.

**Figure 4.**
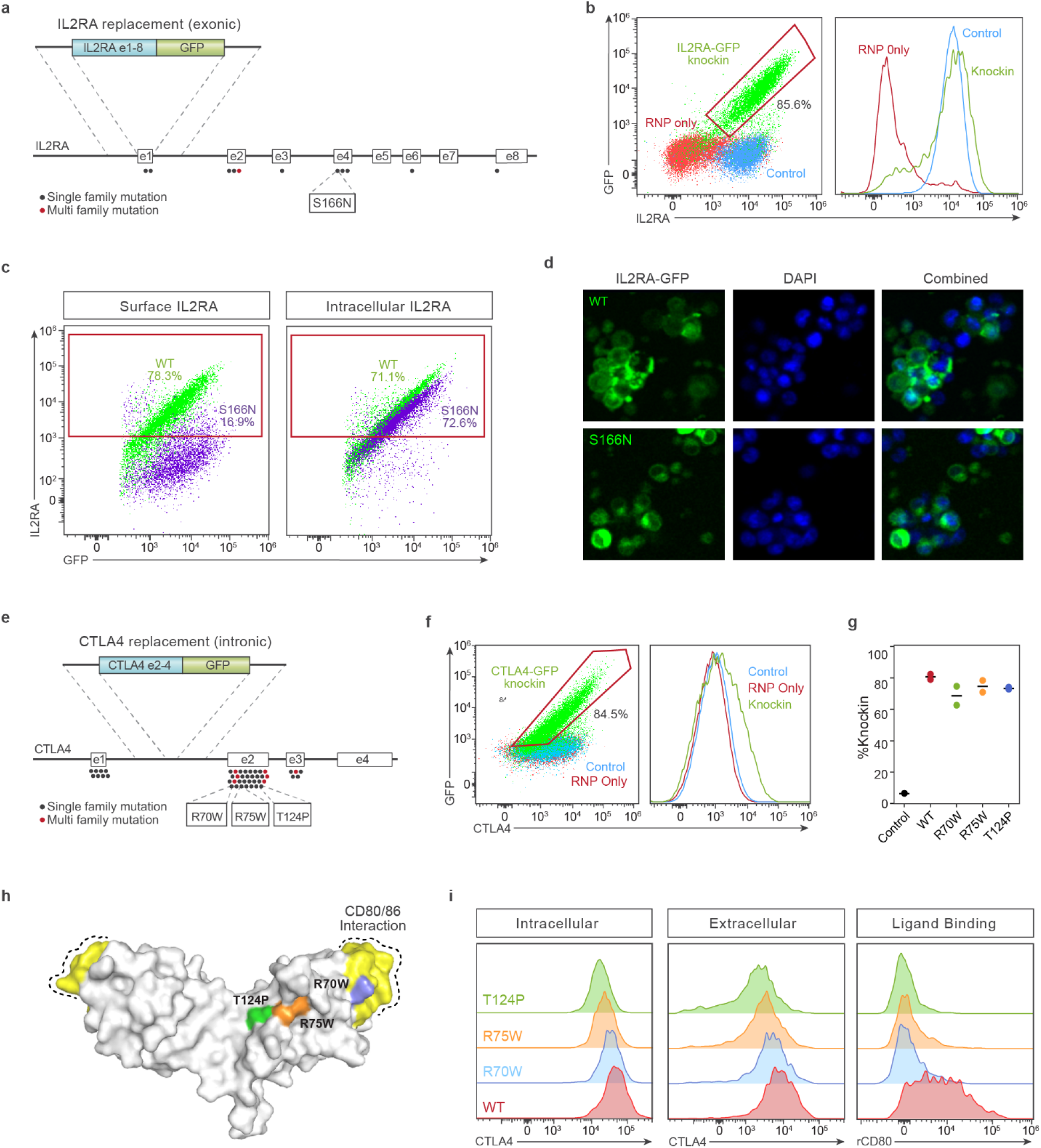
Whole open reading frame (ORF) replacement at target genes for therapeutic and diagnostic human T cell editing. **(a-d)** *IL2RA* exon 1-8 ORF replacement strategy. **(a)** Diagram of the *IL2RA* gene with reported patient coding mutations and knock-in strategy using an IL2RA-GFP fusion protein targeted to exon 1. The S166N mutation examined explored in panel c-d is noted. **(b)** IL2RA and GFP expression in CD4+ T cells electroporated with IL2RA-GFP ssCTS templates and cognate RNP followed by MTX inhibitor combination (green), in comparison to RNP only (red), or no electroporation control cells (blue). **(c)** Comparison of extracellular (surface staining) and intracellular (staining in permeabilized cells which includes total surface and intracellular protein) IL2RA expression with WT and S166N IL2RA-GFP knock-ins assessed by flow cytometry. Percent IL2RA+ is shown for each panel. **(d)** Localization of WT and S166N IL2RA-GFP protein by fluorescence microscopy. **(e-i)** CTLA4 exon 2-4 ORF replacement strategy. **(e)** Diagram of the *CTLA4* gene with reported patient mutations and knock-in strategy using a CTLA4-GFP fusion protein targeted to intron 1. The R70W, R75W, T124P mutations examined in panel g-i are noted. **(f)** CTLA4 and GFP expression in CD4+ T cells electroporated with CTLA4-GFP ssCTS templates and cognate RNP followed by MTX inhibitor combination (green), in comparison to RNP only (red), or no electroporation control cells (blue). **(g)** Quantification of percent knock-in for WT, R70W, R75W, and T124P constructs electroporated with ssCTS templates and treated with the MTX inhibitor combination assess by flow cytometry. **(h)** Structure of CTLA4 dimer with CD80/86 interaction domain highlighted (yellow) along with location of R70W (blue), R75W (orange), and T124P (green) mutations.^35^ **(i)** Comparison of extracellular CTLA4 (surface staining), intracellular CTLA4 (staining in permeabilized cells which includes total surface and intracellular protein), and biotinylated recombinant CD80 ligand interaction stained with Streptavidin-APC in WT, R70W, R75W, and T124P knock-in CD4+ T cells. Each experiment was performed with T cells from 2 independent healthy human blood donors. Error bars indicate standard deviation. RNP = Cas9 Ribonucleoprotein, HDRT = homology-directed-repair template, DAPI = 4’,6-diamidino-2-phenylindole nuclear stain, rCD80 = recombinant CD80.

### Development of a GMP-compatible manufacturing process for non-viral genome engineered T cell therapies

Finally, we sought to generate a clinical-grade process for fully non-viral knock-in of large therapeutic constructs. One of the most immediate applications with demonstrated functional benefit is targeting a CAR insertion to the endogenous *TRAC* locus. This approach greatly enhanced the potency of CD19-specific CAR-T cells in preclinical studies and reduced T cell exhaustion through tightly regulated expression driven by the gene regulatory elements governing normal TCR expression^6^. In contrast to the original rAAV-mediated methods, we adapted this strategy to make use of ssCTS templates targeting the BCMA antigen, a promising target for treatment of Multiple Myeloma that has recently seen FDA-approval for viral CAR-T products (Fig. 1F)^33^. Clinical translation requires transitioning to Good Manufacturing Practice (GMP) compliant reagents, equipment, and processes. For electroporations, we used the Maxcyte GTx platform which provides a GMP-compatible electroporation device with access to FDA Master File along with sterile single-use cuvettes and assemblies that are scalable to the large numbers of cells needed for manufacturing a full patient dose. For genome editing reagents, we used research-grade equivalents that are each available at GMP-grade, including SpyFi Cas9 (a high fidelity Cas9 variant produced at GMP-grade by Aldevron) and chemically synthesized sgRNA also produced at GMP-grade by Synthego^18^. We partnered with Genscript to develop a GMP-compatible process for ssCTS template generation. Encouragingly, Genscript templates encoding a BCMA-CAR knock-in were able to be manufactured at large scale and consistently outperformed our internally generated HDRTs, showing lower levels of toxicity and higher knock-in efficiencies for both ssCTS and dsCTS constructs (Supplementary Fig. 5A).

To demonstrate a large-scale non-viral CAR T manufacturing process, ~100 x 10^6^ primary human T cells were isolated from two healthy donors, activated on Day 0 with CD3/CD28 Dynabeads along with IL-7 and IL-15, electroporated on Day 2 using Maxcyte R-1000 cuvettes, then expanded in G-Rex 100M gas permeable culture vessels to Day 7 or Day 10 (Fig. 5A). Average knock-in efficiencies were 40.4% on Day 7 and 45.8% on Day 10. The final yield of CAR+ cells was >5 x 10^8^ by Day 7 and >1.5 x 10^9^ by Day 10 for both donors, well within the range needed to generate a full patient dose of ~100 x 10^6^ CAR+ cells (Fig. 5B-D). While the addition of small molecule inhibitors improved knock-in efficiencies to >60%, we observed a reduction in live cell counts such that the final yield of CAR+ cells were decreased in comparison to ssCTS templates alone (Fig. 5B-D, Supplementary Fig. 5C-D). The majority of CAR+ cells demonstrated an immunophenotype consistent with a T stem cell memory (Tscm) population on day 10 of expansion based on CD45RA/CD62L expression and confirmed with additional markers as CD45RA^+^CD62L^+^CD45RO^-^CCR7^+^CD95^+^ (Fig. 5E, Supplementary Fig. 5B-C). *In vitro* assays demonstrated efficient killing of BCMA+ MM1S myeloma cell lines in contrast to unmodified T cells expanded from the same donors (Fig. 5F). Altogether, these results demonstrate a fully non-viral manufacturing process capable of high efficiency T cell engineering at clinical scale which may be transitioned to full-GMP manufacturing and quickly adapted toward additional targets.

**Figure 5.**
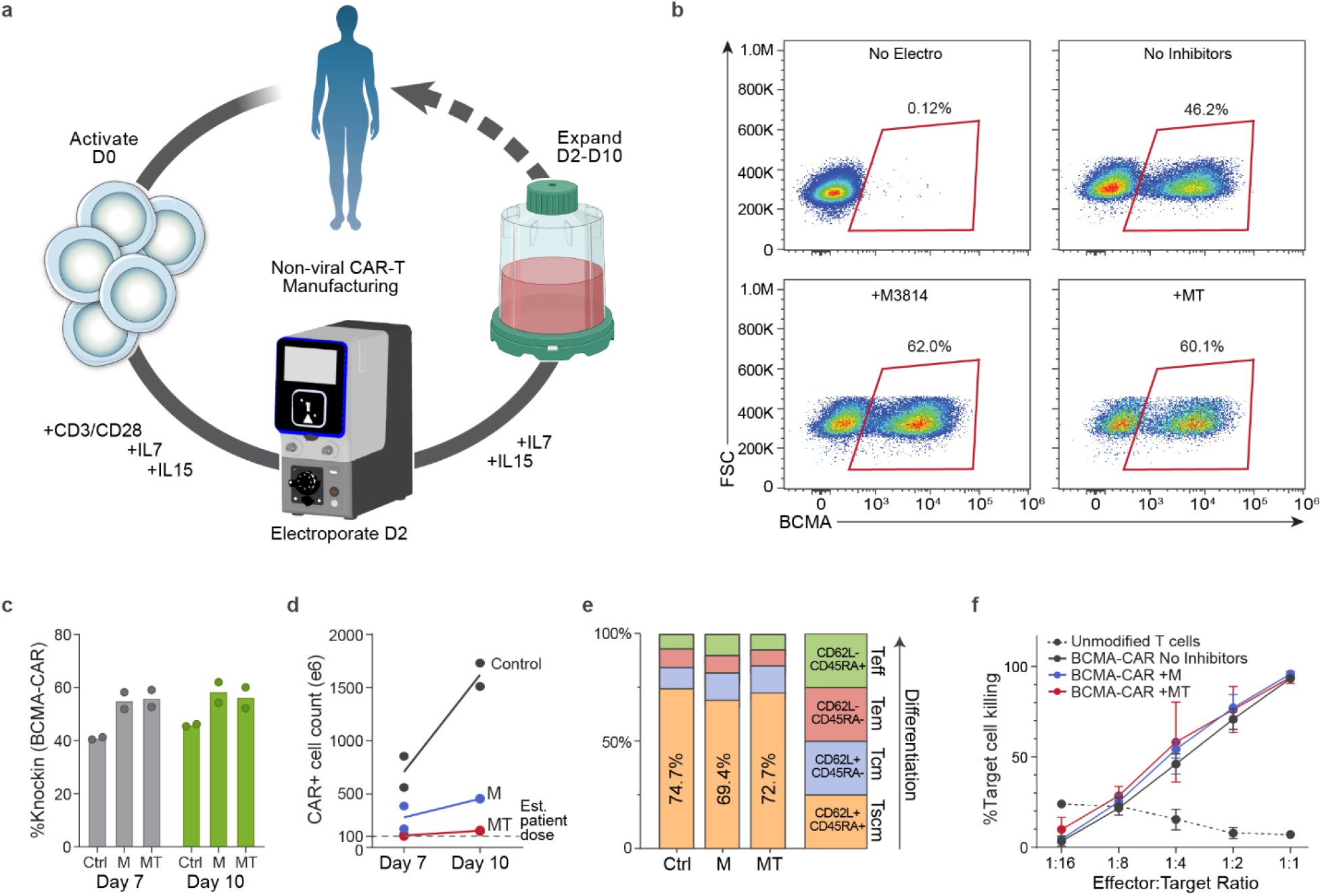
GMP-compatible process for non-viral CAR-T cell manufacturing. **(a)** Diagram of non-viral CAR-T cell manufacturing process. T cells are isolated from peripheral blood and activated on Day 0 with anti-CD3/anti-CD28 Dynabeads, IL-7, and IL-15. Cells are electroporated using the Maxcyte GTx electroporator on Day 2 with Cas9 RNPs + ssCTS HDRTs and then expanded for an additional 7-10 days using G-Rex 100M culture vessels supplemented with IL-7 + IL-15. **(b)** Representative Day 10 flow plots showing BCMA-CAR knock-in for Control (No inhibitors), M3814, and M3814 + TSA (MT) conditions. **(c)** BCMA-CAR knock-in rates on Day 7 and Day 10 for each condition. **(d)** Absolute number of CAR+ cells on Day 7 and Day 10. Dotted line shows an estimated patient dose of ~100 x 10^6^ CAR+ T cells. **(e)** T cell immunophenotype on Day 10 based on CD45RA and CD62L expression. **(f)** *In vitro* killing of BCMA+ MM1S multiple myeloma cell lines in comparison to unmodified T cells from same blood donors. Each experiment was performed with T cells from 2 independent healthy human blood donors. Error bars indicate standard deviation. Panel a was generated in part using graphics created by Biorender.com. RNP = Ribonucleoprotein, CTS = Cas9 Target Site, ssCTS = ssDNA HDRT + CTS sites, HDRT = homology-directed-repair template, M = M3814, MT = M3814 + TSA, Tscm = T stem cell memory, Tcm = T central memory, Tm = T effector memory, Teff = T effector.

## Discussion

The ability of CRISPR genome engineering to introduce targeted sequence replacements or insertions in primary human cells holds immense promise for studying disease variants, correction of genetic diseases, and reprogramming cellular functions for the next-generation of cell-based therapeutics. Here we report advances that improve HDR efficiency and yield with large non-viral ssCTS templates and small molecule inhibitor combinations. We apply this technology across diverse genetic loci, knock-in constructs, and primary hematopoietic cell types, demonstrating their utility for the generation of universal gene correction strategies, disease variant modeling, and GMP-compatible manufacturing processes.

ssCTS hybrid repair templates – alone or in combination with small molecule inhibitors – provide a broadly useful tool to promote CRISPR-based HDR. The technology reported here demonstrated >7-fold increases at some sites. However, by testing knock-in across a broad array of target sites, we did observe variation even with different RNPs targeting the same gene. Variable knock-in rates and toxicity could be affected by unique features of the target site (or off-target effects) at the local sequence or epigenetic level. Recent work has highlighted that some gRNA targets exhibit distinct repair pathway preferences^21^. A detailed analysis of repair outcomes at the sequence level, reliance on alternative repair pathways, and evaluation of off-target effects may help identify the source of this variability and inform future design of genome targeting strategies.

The relatively high purity and high yield of live cells achieved here with large genome replacements provides a powerful tool to probe DNA sequence function in primary human cells. We can now directly test the function of individual coding or non-coding genome sequences for mechanistic studies or to confirm the clinical relevance of disease variants. The most recent classification of Inborn Errors of Immunity (aka Primary Immunodeficiencies or PID) from the 2019 International Union of Immunological Societies (IUIS) update identifies >400 monogenic immune disorders with 65 new genes implicated since 2017^23^. Families with these diseases demonstrate a spectrum of mutations scattered throughout these genes and interpretation of novel variants of unknown significance (VUS) is a persistent challenge to diagnosis and appropriate patient management. Routine interrogation of these VUS at endogenous loci within the relevant primary human cell type may now be feasible. Here we demonstrate application of our non-viral approaches at a variety of PID-associated genes, in some cases achieving knock-in efficiencies >80% without selection and allowing us to evaluate the functional consequences of disease-causing mutations within primary human T cells. We further demonstrate the ability to extend these approaches to alternative hematopoietic cell types and large knock-in pools, providing a foundation for high-throughput functional screens that may be used to examine the immense variety of PID-associated genetic variants.

Enhanced CRISPR-based genome targeting with ssCTS templates also provides opportunities to re-write sequences in primary somatic cells to treat patients. Here we show the potential of non-viral approaches to generate high-efficiency universal open reading frame (ORF) replacements for two genes, *IL2RA* and *CTLA4*, both associated with severe autoimmunity and immune dysfunction affecting primary human T cells. Flexible, non-viral gene replacement strategies – in hematopoietic stem and progenitor cells or in more terminally differentiated cell types such as T cells – could give more patients access to curative cellular therapies. More broadly, the ability to efficiently knock-in large sequences into specified genome locations opens the door to synthetic reprogramming to generate powerful cellular medicines. Here we demonstrate clinical-scale, non-viral manufacturing of T cells engineered to have chimeric antigen receptors (CARS) expressed under the gene regulatory control of endogenous *TCR-alpha*, which has been reported to have favorable properties. Eventually, this process should support robust manufacturing of even more complex synthetic gene programs integrated into targeted genome sites to drive potent cell therapy functions for diverse, complex human diseases.

Altogether, we have developed a variety of tools and applications that markedly improve non-viral genome editing and demonstrate the power of these methods to correct, modify, and reprogram primary human cells. We have applied these approaches predominantly toward genome targets relevant for human T cell editing, demonstrating applications for functional genetic screens or therapeutic genome engineering. However, we also show the feasibility of applying ssCTS templates to a range of relevant human cell types and these approaches may be extended for many alternative applications, including targeting the >400 genes associated with a PID or incorporation of a wide variety of novel synthetic biology constructs. These studies demonstrate the capacity of fully non-viral HDR to mediate complex and targeted genome modifications with high efficiency and yield, setting the stage for a number of research, diagnostic, and manufacturing applications which we hope will reduce the complexity of clinical translation and streamline the development of new therapies.

## Methods

### Cell Culture

Primary adult blood cells were obtained from anonymous healthy human donors as a leukapheresis pack purchased from StemCell Technologies, Inc. or Allcells Inc, or as a Trima residual from Vitalant. If needed, peripheral blood mononuclear cells were isolated by Ficoll-Paque (GE Healthcare) centrifugation. Primary human cell types were then further isolated by positive and/or negative selection using EasySep magnetic cell isolation kits purchased from StemCell for CD3+ T cells (Cat #17951), CD4+ T cells (Cat #17952), CD8+ T cells (Cat #17953), B cells (Cat #17954), NK cells (Cat #17955), or CD4+CD127lowCD25+ regulatory T cells (Cat #18063) per manufacturer instructions. Primary human γδ T cells were isolated using a custom γδ T cell negative isolation kit without CD16 and CD25 depletion obtained from StemCell. Primary adult peripheral blood G-CSF-mobilized CD34+ hematopoietic stem cells were purchased from StemExpress, LLC.

With the exception of GMP-compatible scale-up experiments (described separately below), isolated CD3+, CD4+, CD8+, and γδ T cells were activated at 1 x 10^6^ cells mL^-1^ for 2 days in complete XVivo15 medium (Lonza) (5% fetal bovine serum, 50 μM 2-mercaptoethanol, 10 mM N-acetyl L-cysteine) supplemented with anti-human CD3/CD28 magnetic Dynabeads (CTS, ThermoFisher) in a 1:1 ratio with cells, 500 U mL^-1^ of IL-2 (UCSF Pharmacy), and 5 ng mL^-1^ of IL-7 and IL-15 (R&D Systems). Regulatory T cells were activated at 1 x 10^6^ cells mL^-1^ for 2 days in complete XVivo15 supplemented with magnetic Treg Xpander CTS Dynabeads (ThermoFisher) at a 1:1 bead to cell ratio and 500 U mL^-1^ of IL-2 (UCSF Pharmacy). Isolated B cells were activated at 1 x 10^6^ cells mL^-1^ for 2 days in IMDM medium (ThermoFisher) with 10% fetal bovine serum, 50 μM 2-mercaptoethanol, 100 ng mL^-1^ MEGACD40L (Enzo), 200 ng mL^-1^ anti-human RP105 (Biolegend), 500 U mL^-1^ IL-2 (UCSF Pharmacy), 50 ng mL^-1^ IL-10 (ThermoFisher), and 10 ng mL^-1^ IL-15 (R&D Systems). Isolated NK cells were activated at 1 x 10^6^ cells mL^-1^ for 5 days in XVivo15 medium (Lonza) with 5% fetal bovine serum, 50 μM 2-mercaptoethanol, 10 mM N-acetyl L-cysteine, 1000 U mL^-1^ IL-2, and MACSiBead Particles pre-coated with anti-human CD335 (NKp46) and CD2 antibodies based on manufacturer guidelines (Miltenyi Biotec). Primary adult CD34+ HSCs were cultured at 0.5 x 10^6^ cells per mL in SFEMII medium supplemented with CC110 cytokine cocktail (StemCell).

For GMP-compatible scale-up experiments, CD3+ T cells were activated with anti-human CD3/CD28 magnetic Dynabeads (CTS, ThermoFisher) in a 1:1 ratio with 100 U mL^-1^ of IL-7 and 10U mL^-1^ IL-15 (R&D Systems) in tissue culture flasks. Post-electroporation, cells were expanded in G-Rex 100M gas-permeable culture vessels (Wilson Wolf) supplemented with 100 U mL^-1^ of IL-7 and 10U mL^-1^ IL-15 every 2-3 days for a total 7 or 10 day expansion as indicated.

### RNP Formulation

For most experiments (excluding GMP-compatible scale-up described separately below), ribonucleoproteins (RNP) were produced by complexing a two-component gRNA to Cas9 with addition of either a Poly-glutamic acid (PGA) or ssDNAenh electroporation enhancer, as previously described^11^. Synthetic CRISPR RNA (crRNA, with guide sequences listed in Supplementary Table 1) and trans-activating crRNA (tracrRNA) were chemically synthesized (Edit-R, Dharmacon Horizon), resuspended in 10 mM Tris-HCl (pH 7.4) with 150 mM KCl or IDT duplex buffer at a concentration of 160 μM, and stored in aliquots at −80 °C. The ssDNAenh electroporation enhancer was purchased was synthesized by IDT (with sequence listed in Supplementary Table 2), resuspended to 100 μM in water, and stored at −80 °C. 15–50 kDa PGA was purchased from Sigma and resuspended to 100 mg ml-1 in water, sterile filtered, and stored at −80 °C prior to use.

To make gRNA, aliquots of crRNA and tracrRNA were thawed, mixed 1:1 v/v, and annealed by incubation at 37 °C for 30 min to form an 80 μM gRNA solution. PGA or ssDNAenh were mixed into gRNA solutions at a 0.8:1 volume ratio prior to adding 40 μM Cas9-NLS (Berkeley QB3 MacroLab) at a 1:1 v/v to attain a molar ratio of sgRNA:Cas9 of 2:1. Final RNP mixtures were incubated at 37°C for 15-30 minutes after a thorough mix. Based on a Cas9 protein basis, 50 pmol of RNP was used for each electroporation.

For GMP-compatible scale-up experiments, synthetic single guide RNA (sgRNA) was purchased from Synthego, resuspended to 160 μM, aliquoted and stored at −80°C. SpyFi Cas9 nuclease was purchased from Aldevron LLC, aliquoted, and stored at −20°C. For RNP formulation, aliquots of ssDNAenh and sgRNA solutions were thawed and mixed at a 0.8:1 volume ratio prior to adding SpyFi Cas9 at a 2:1 molar ratio of sgRNA:Cas9. Final RNP mixtures were incubated at 37°C for 15–30 minutes prior to electroporation.

### HDRT Template Preparation

Short ssDNA HDRTs (<200bp) were directly synthesized (Ultramer oligonucleotides, IDT), resuspended to 100 uM in dH20, and stored at −20 °C prior to use. Long dsDNA HDRTs encoding various gene insertions (see Supplementary Table 1) and 300–600 bp homology arms were synthesized as gBlocks (IDT) and cloned into a pUC19 plasmid inhouse or purchased directly from Genscript Biotech. These plasmids then served as a template for generating a PCR amplicon. CTS sites were incorporated through additional 5’ sequence added to the base PCR primers (see Supplementary Table 1 for sequences). Amplicons were generated with KAPA HiFi polymerase (Kapa Biosystems), purified by SPRI bead cleanup, and resuspended in water to 0.5–2 μg μl–1 measured by light absorbance on a NanoDrop spectrophotometer (Thermo Fisher), as previously described^11,12^.

For most experiments requiring long ssDNA (excluding GMP-compatible scale-up described separately below), a ssDNA isolation protocol adapted from Wakimoto et al. using biotinylated primers and streptavidin-coated magnetic beads was used^34^. Amplicons were generated as described above using primers that include a 5’ biotin modification (IDT) on either the forward or reverse PCR primer. ~20uL Streptavidin C1 Dynabeads (ThermoFisher, Cat # 65001) per 1 picomole of amplicon were rinsed 3 times with 1X Binding & Wash (B&W) buffer (prepared at 2X concentration and stored at RT using 10mL 1M TRIS-HCl pH 7.5, 2mL 0.5M EDTA, 116.88g NaCl, 1L dH20) using magnetic separation. The washed beads and the PCR amplicon were then resuspended in B&W buffer for 30 minutes at room temperature to capture the biotinylated DNA. The mixtures were washed twice with B&W buffer after which the supernatant was removed and replaced with 0.125M NaOH Melt Solution (prepared fresh) to denature the dsDNA. The solution is placed back on the magnet for 5 minutes and the supernatant containing the non-biotinylated strand is removed gently with non-stick pipettes and mixed immediately with Neutralization Buffer (100 uL 3M Sodium Acetate pH 5.2 and 4.9 mL 1X TE Buffer, prepared fresh). Resulting ssDNA was purified and concentrated using a SPRI bead cleanup, as described previously, and quantified on a Nanodrop spectrophotometer (Thermo Fisher).

### Large-Scale ssDNA Production

For GMP-compatible scale-up experiments, research grade long single-stranded DNA was manufactured at large scale by Genscript Biotech via a proprietary isothermal enzymatic reaction process (PCT/CN2019/128948). To be brief, sequence verified template on plasmid vector is first be converted into uridine modified linear dsDNA fragments via PCR amplification. The linear dsDNA is then treated with USER^®^ Enzyme and T4 ligase (Cat. #M5505S and M0202T, New England BioLabs) to form a self-ligated dsDNA circle with nicking sites. This nick containing dsDNA circle is used as an amplification template for rolling circle amplification, which is carried out by phi29 DNA polymerase (Cat. # M0269L, New England BioLabs) in a high fidelity and linear amplification manner. The product of rolling circle amplification is ssDNA concatemers with repeats of target fragment and a palindromic adapter sequence. The annealing process is followed to let the palindromic adapter sequence form a hairpin structure, and then BspQI restriction enzyme (Cat. # R0712L, New England BioLabs) is added in the reaction system to recognize the stem part of the hairpin and digest the concatemer intermediates into target ssDNA monomers and hairpin adapters. The crude product is further purified by EndoFree^®^ Plasmid Maxi Kit (Qiagen, Cat. # 12362), to harvest the target ssDNA and remove hairpin adapters, enzymes, reaction buffer, and endotoxin residues.

For production of the 2,923nt BCMA-CAR encoding ssDNA material, amplification primers were synthesized to add specially designed adapter sequences at the 5’ and 3’ ends of the target sequence via PCR method. The uridine modified forward and reverse primer sequences manufactured by Genscript were: 5’-AACTATACUACGTCAATCGGCTCTTCACACTACTACAGTGCCAATAG-3’ and 5’-TATAGTUACGTCAATCGGC TCTTCACACCGTCTGACTAACATAACCTG-3’, respectively. The cycle number of the PCR reaction was set as 20, and 300 μg of linear dsDNA fragment was produced and purified by QIAquick PCR Purification Kit (Qiagen, Cat. # 28706). All of the purified 300μg of linear dsDNA was treated with USER enzyme and T4 ligase to prepare the rolling circle amplification template, and then, it was used as the template for a 100 mL RCA reaction. All of the isothermal enzymatic reactions and annealing process were done on Eppendorf ThermoMixer^®^ C. The final purified ssDNA sample was eluted with nuclease-free water (Sigma Aldrich, Cat. # W4502) from the silica column of an EndoFree^®^ Plasmid Maxi Kit, and then passed single-use 0.22 μm sterile filter (Millipore, Cat. # SLGV033RS). Before lyophilization and final packaging, the ssDNA material was quantified by Nanodrop One^C^ (Thermo Fisher) by UV 260 nm absorbance in single-stranded DNA mode. The sequence integrity was confirmed by Sanger sequencing, and the homogeneity was measured by 2% agarose gel electrophoresis as a single band. Quality control for biosafety of the ssDNA material was also evaluated: endotoxin residue was determined as ≤ 10 EU/mg by an endotoxin test kit (Bioendo, Cat. # KC5028), protein residue level was below the minimum detection threshold of Micro BCA Protein Assay Kit (Thermo Fisher, Cat. # 23235), and no bacterial colonies formed in bioburden detection.

### Electroporation and use of small molecule inhibitors

Except for GMP-compatible scale-up experiments, primary cells were isolated on day 2 of culture (HSCs, CD3+, CD4+, CD8+, γδ, and regulatory T cells) or day 5 (NK cells) and electroporated using the Lonza 4D 96-well electroporation system as previously described^11^. CD3+, CD4+, CD8+, γδ, and regulatory T cells were debeaded using an EasySep magnet (StemCell). Immediately prior to electroporation, cells were centrifuged at 90g for 10 minutes and then resuspended at 0.4 x 10^6^ HSCs, 0.5 x 10^6^-1.0 x 10^6^ T cells, 0.5 x 10^6^ NK cells, or 0.5 x 10^6^ B cells per 20uL Lonza P3 buffer. HDRT and RNP formulations were mixed and incubated for at least 5 minutes, then combined with cells and transferred to the Lonza 96-well electroporation shuttle. B cells, NK cells, and all T cell subtypes were electroporated using pulse code EH-115 while HSCs were electroporated with pulse code ER-100. Following electroporation, cells were rescued with prewarmed growth media and incubated for at least 15 minutes. Cells were then transferred to fresh plates or flasks and diluted to 0.5-1.0 x 10^6^ cells mL^-1^ in each respective growth medium as described above. Fresh cytokines and media were added every 2-3 days.

Trichostatin A (TSA) (Cayman Chemical), Nedisertib (M3814) (MedKoo Biosciences), XL413 hydrochloride (XL413) (Fisher Scientific), NU7441 (Fisher Scientific), and Alt-R HDR enhancer (IDT) were prepared and stored as aliquots per manufacturer guidelines. For experiments using small molecule inhibitors, cells were incubated with the indicated concentrations upon addition of fresh growth media following the 15 minute rescue step, and removed by media exchange after 24 hours. Longer incubation times of 48 and 72 hours did not improve knock-in efficiency further and were associated with increased toxicity (data not shown).

For GMP-compatible scale-up experiments, activated cells were separated from beads on day 2 and centrifuged for 10 minutes at 90g. After removing the supernatant, cells were resuspended in Maxcyte Electroporation Buffer at 200 x 10^6^ cells mL^-1^. HDRTs and RNPs were mixed and incubated for at least 5 minutes before being combining with cells. The mixture was then transferred to Maxcyte OC-1000 electroporation cuvettes. Cuvettes were filled up to ~60% of the total volume (~600uL) and electroporated with pulse code Expanded T cell 4-2. Immediately following electroporations, ~400uL prewarmed XVivo15 media was added to the cuvette and cells were incubated for 15 minutes, then transferred to G-Rex culture vessels as described above.

### Flow Cytometry

All flow cytometry was performed on an Attune NxT flow cytometer with a 96-well autosampler (Thermo Fisher Scientific). Unless otherwise indicated, cells were collected 3-5 days post-electroporation, resuspended in FACS buffer (1%–2% BSA in PBS) and stained with Ghost Dye red 780 (Tonbo) and the indicated cell-surface and intracellular markers (see Supplementary Table 2 for antibodies). For intracellular staining, cells were stained for surface markers and then prepared for intracellular staining using True-Nuclear Transcription Factor staining kits (Biolegend). For experiments demonstrating stimulation response, cells were re-activated 24 hours prior to analysis using ImmunoCult Human CD3/CD28/CD2 T Cell Activation reagent (StemCell). Analysis was done using FlowJo v10 software. All gating strategies included exclusion of subcellular debris, singlet gating, and live:dead stain. Final graphs were produced with Prism (GraphPad), and figures were compiled with Illustrator (Adobe).

---

**Supplementary Figure 1.**
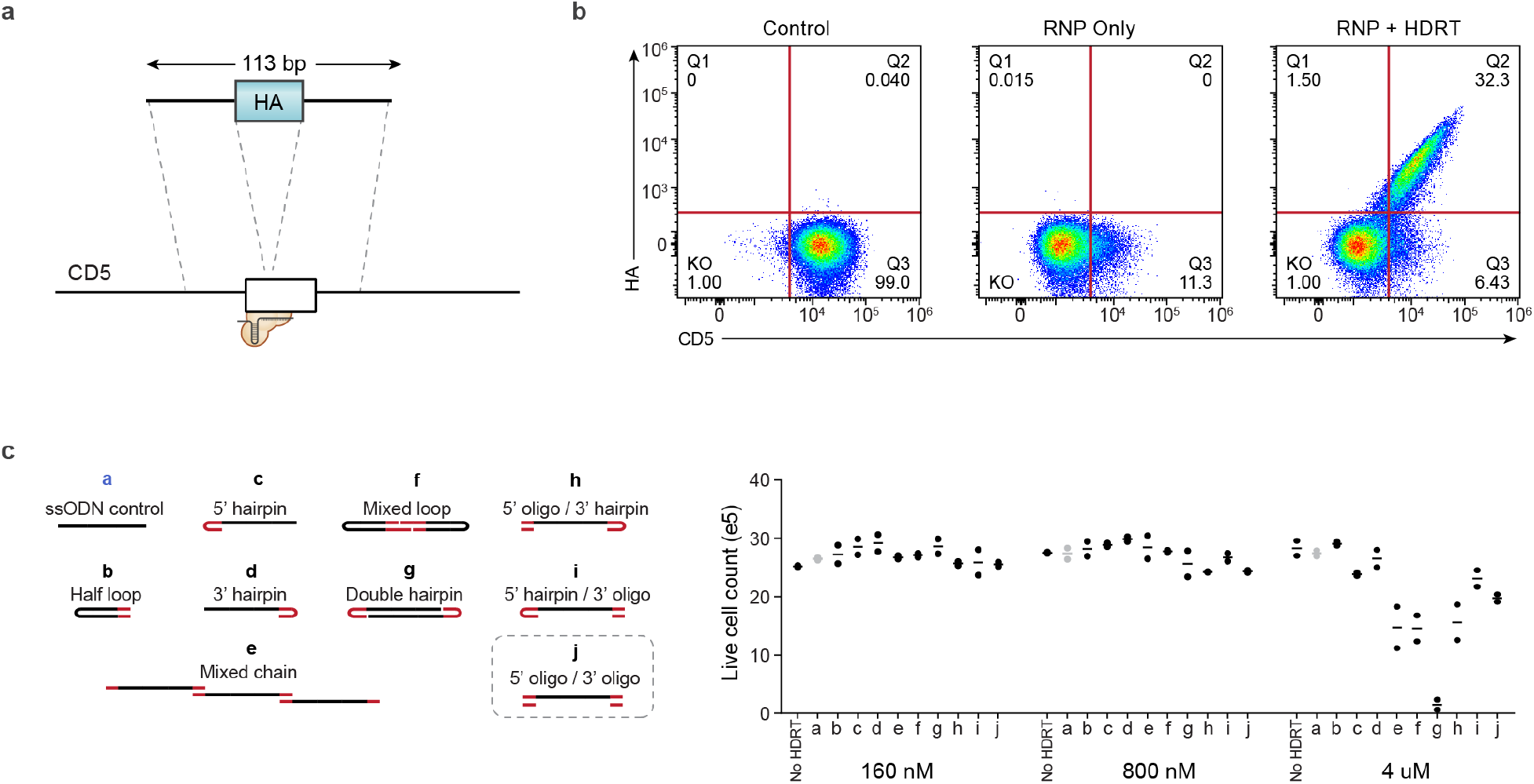
Screening with short CD5-HA ssCTS templates. **(a)** Diagram of CD5-HA knock-in strategy and control ssDNA HDRTs. **(b)** Representative flow cytometry plots demonstrating CD5-HA knock-in. **(c)** Live cell counts for each ssCTS design using a CD5-HA knock-in construct at 160 nM – 4uM concentration. Each experiment was performed with T cells from 2 independent healthy human blood donors. Error bars indicate standard deviation. RNP = Ribonucleoprotein, HDRT = homology-directed-repair template.

**Supplementary Figure 2.**
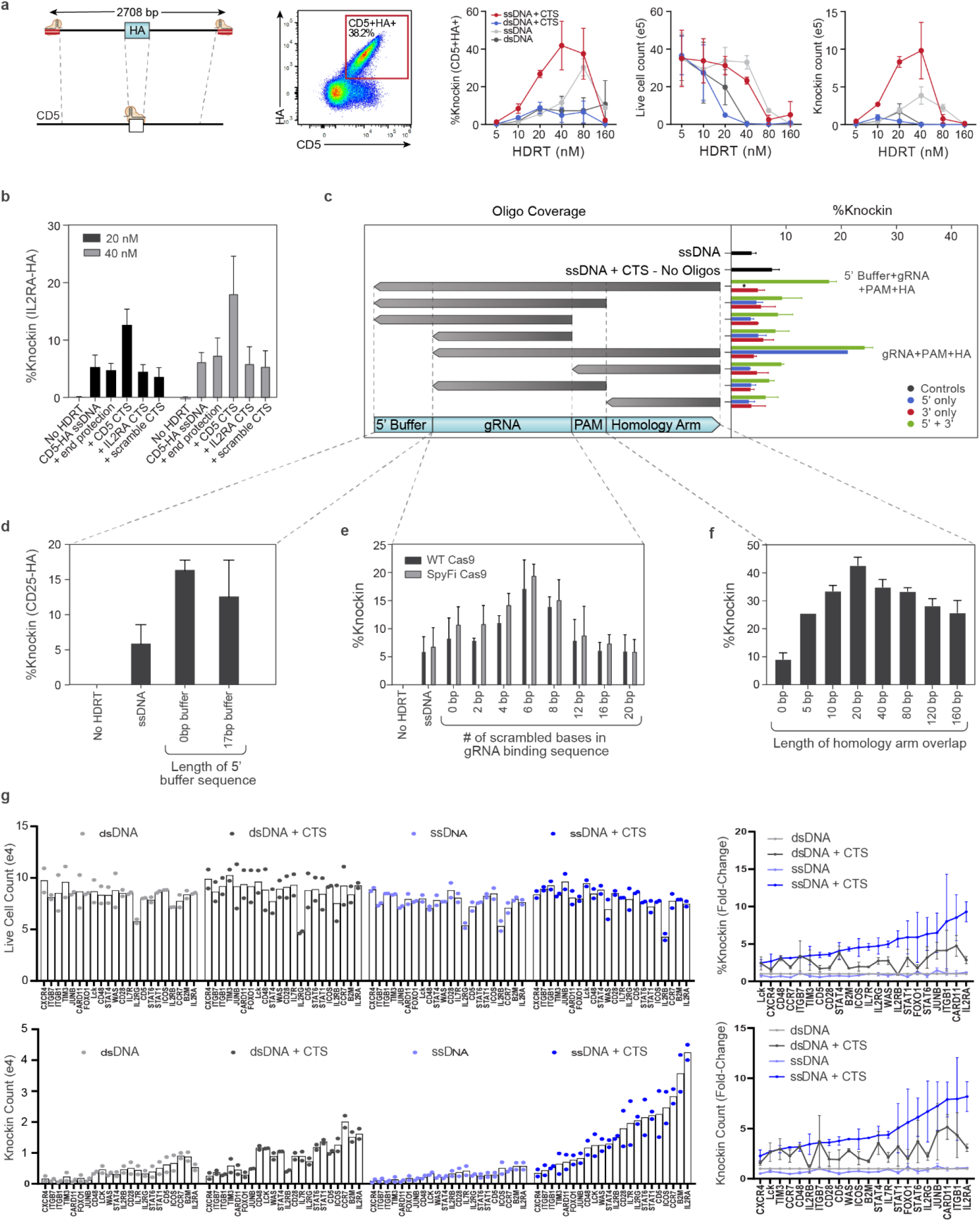
Optimization of ssCTS design with large CD5-HA HDRTs. **(a-f)** Comparison of different CTS designs with a large ~2.7kb CD5-HA knock-in construct. **(a)** Diagram of long CD5-HA knock-in strategy, representative flow cytometry plot, %knock-in, live cell counts, and knock-in cell yield counts. **(b)** Comparison of CTS with a gRNA target sequence that is specific for the cognate RNP (+ CD5 CTS), an alternative gRNA sequence (+ IL2RA CTS), a CTS incorporating a PAM site and scrambled gRNA sequence (+ scramble CTS), or an equivalent amount of dsDNA within the 5’ end of the homology arm (+ end protection). **(c)** Comparison of complementary oligos covering different regions of the CTS and surrounding sequences. Constructs with CTS sites on both 5’ and 3’ end (green bars), 5’ end only (blue blurs), or 3’ end only (red bars) are shown on the right panel. **(d)** Evaluation of varied 5’ ends including different length of buffer sequence upstream of the CTS site. *indicates no data available for the marked column. **(e)** Comparison of CTS with different numbers of scrambled bases at the 5’ end of the gRNA target sequence using WT or SpyFi Cas9. **(f)** Length of homology arm that is covered by the complementary oligonucleotide. **(g)** Comparison of HDRT variations for knock-in constructs targeting a tNGFR marker across 22 different target loci. Shown for each construct are live cell counts, knock-in cell count yields, relative %knock-in and relative knock-in counts compared to dsDNA templates. Each experiment was performed with T cells from 2 independent healthy human blood donors. Error bars indicate standard deviation. CTS = Cas9 Target Site, PAM = Protospacer Adjacent Motif, HDRT = homology-directed-repair template

**Supplementary Figure 3.**
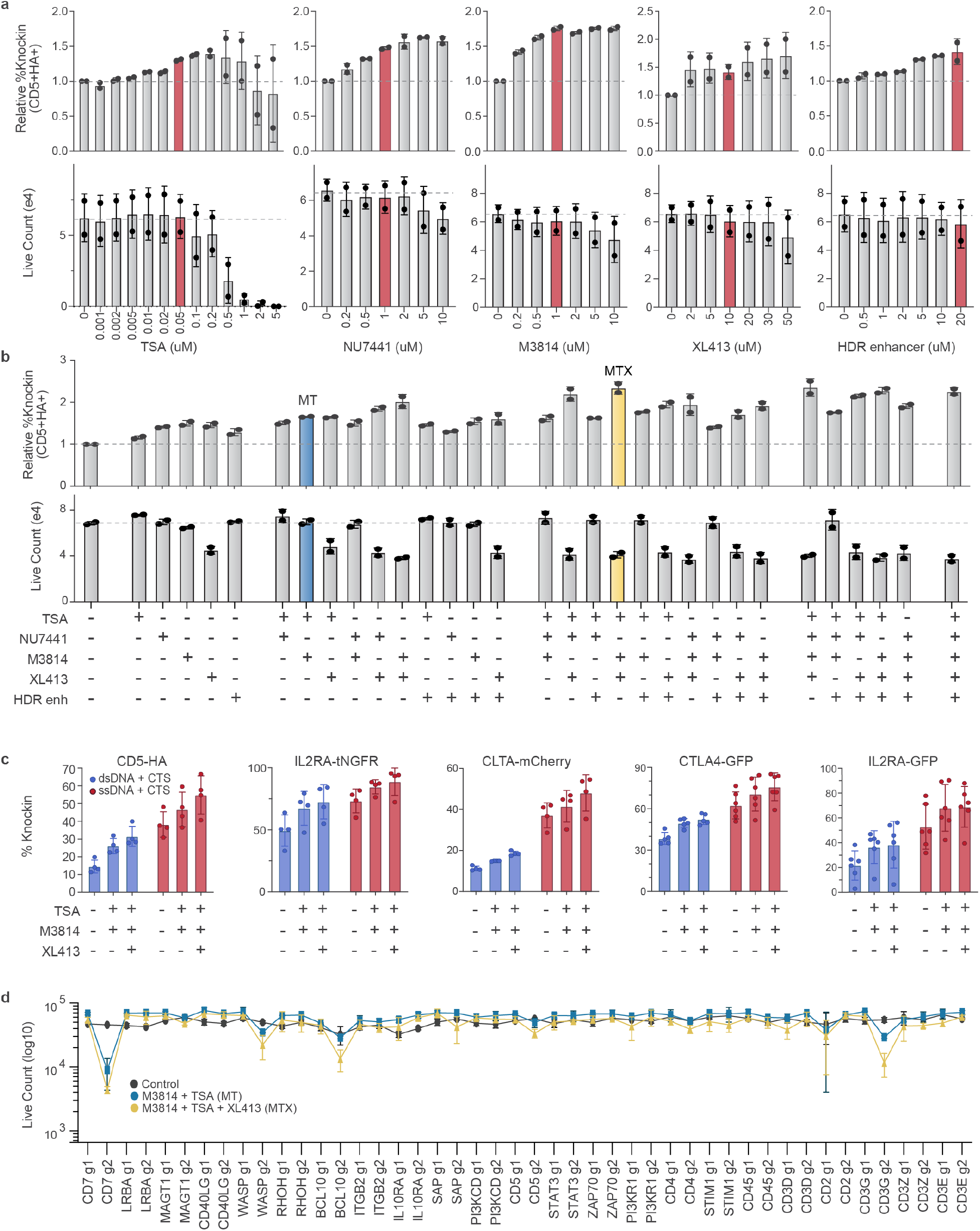
Evaluation of small molecule inhibitors to boost knock-in in primary human T cells. **(a)** Evaluation of relative increase in %knock-in using an ssDNA CD5-HA knock-in construct over varied concentrations of 5 different small molecule inhibitors assessed by flow cytometry. Red bars indicate concentrations chosen for subsequent experiments. **(b)** Comparison of relative %knock-in (top) and live cell counts (bottom) with small molecule inhibitor combinations. Combinations chosen for subsequent experiments are highlighted in blue (MT) and yellow (MTX). **(c)** Comparison of dsCTS and ssCTS templates in combination with small molecule inhibitors for 5 different knock-in constructs including a large CD5-HA HDRT (~2.7kb), a tNGFR knock-in to the *IL2RA* gene (~1.5kb), an mCherry fusion in the Clathrin gene (~1.5kb), a near full length CTLA4-GFP fusion to the *CTLA4* gene (~2.1kb), and a full length IL2RA-GFP fusion to the *IL2RA* gene (~2.3kb). **(d)** Evaluation of live cell counts with MT and MTX inhibitor combinations using 44 different knock-in constructs targeting a tNGFR marker across 22 different target loci with 2 gRNA per gene (g1 and g2). Panel a, b, and d were each performed with T cells from 2 independent healthy human blood donors. Panel c was performed with T cells from 4-6 independent healthy human blood donors. Error bars indicate standard deviation. CTS = Cas9 Target Site, HDRT = homology-directed-repair template, dsCTS = dsDNA + CTS HDRT, ssCTS = ssDNA + CTS HDRT, TSA = Trichostatin A, HDR Enhancer = IDT Alt-R HDR Enhancer, MT = M3814 + TSA, MTX = M3814 + TSA + XL413.

**Supplementary Figure 4.**
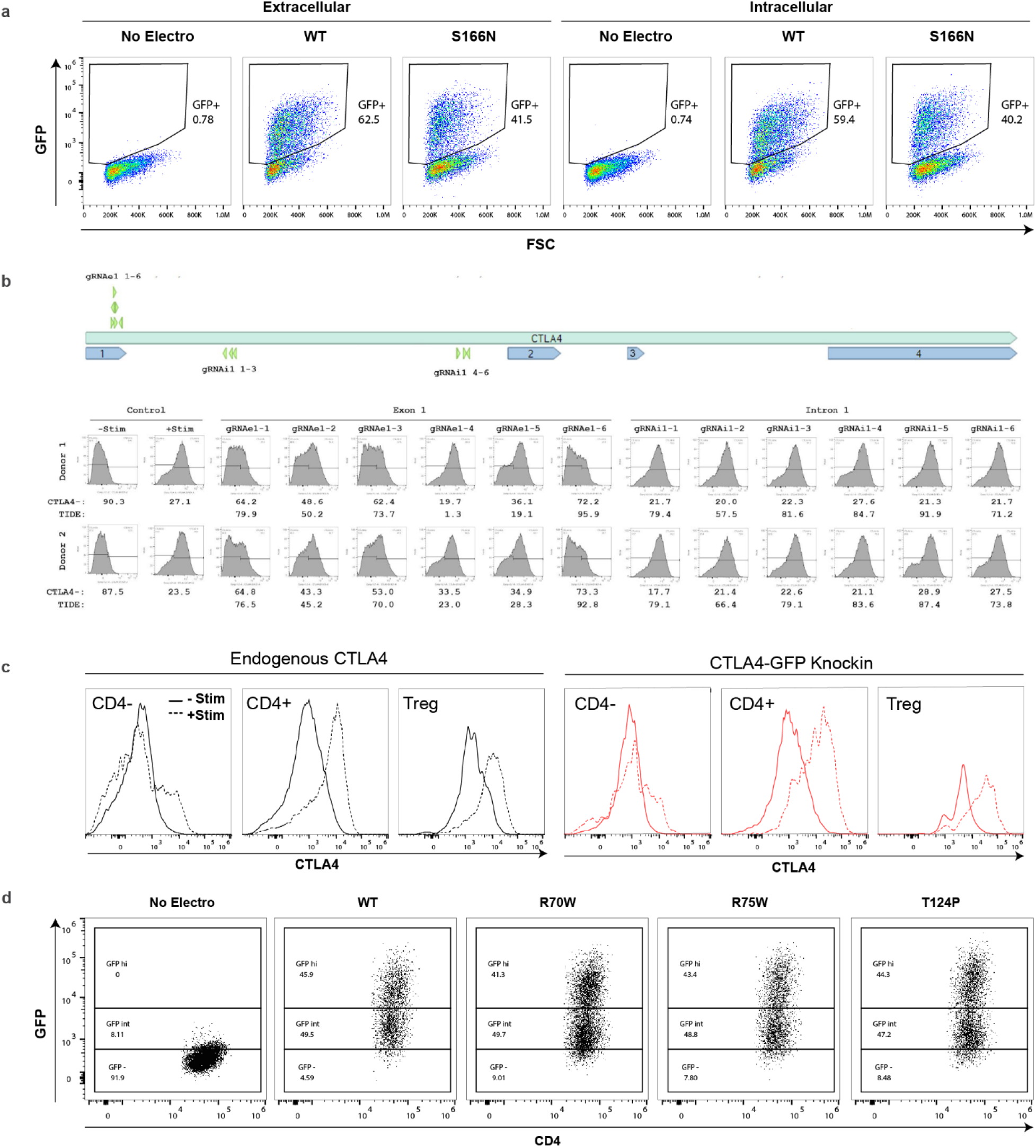
*IL2RA* and *CTLA4* ORF replacement strategies. **(a)** Gating for GFP+ cells is shown with WT and S166N IL2RA-GFP knock-in constructs. **(b)** Diagram of the *CTLA4* gene (top), CTLA4 protein levels (bottom), and cutting efficiency (bottom) illustrating a screening panel of 12 gRNAs examined within exon 1 and intron 1. gRNAs were assessed in activated CD4+ T cells for protein disruption by CTLA4 flow cytometric analysis (flow plots and top row of numbers demonstrate the % of CTLA4-negative cells for each donor), and for cutting efficiency as determined by TIDE indel analysis^36^ (bottom row of numbers indicate the %indel at target locus). **(c)** CTLA4 expression levels assessed by flow cytometry with endogenous protein (black) and WT CTLA4-GFP knock-in protein (red) are shown for CD4-T cells, CD4+ T cells, and regulatory T cells with (dotted line) and without (solid line) stimulation. **(d)** Gating for GFPhi cells is shown for WT, R70W, R75W, and T124P CTLA4-GFP knock-in cells. Each experiment was performed with T cells from 2 independent healthy human blood donors. Error bars indicate standard deviation. WT = Wild-Type, Treg = regulatory T cell.

**Supplementary Figure 5.**
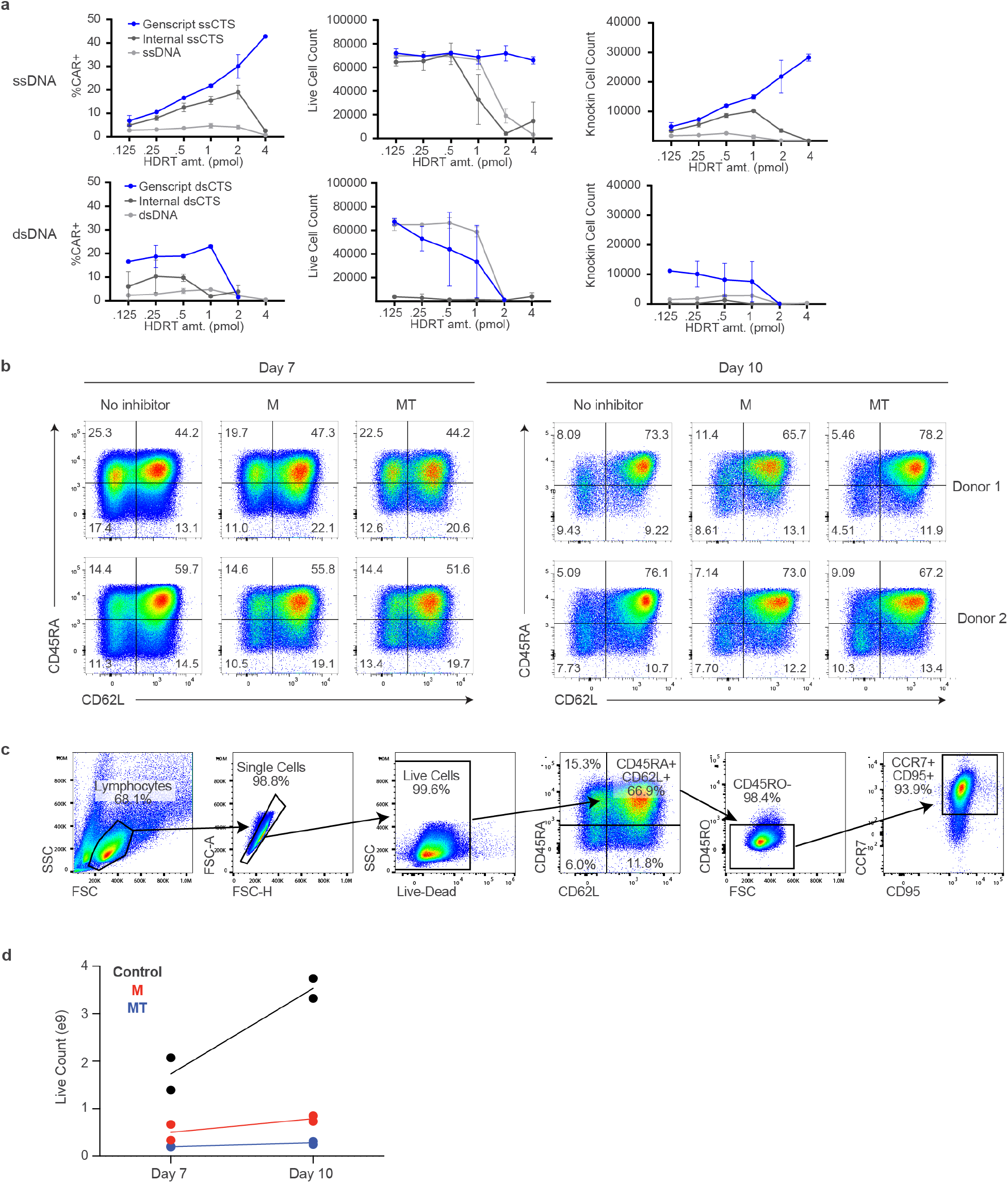
Non-viral CAR-T development with GMP-compatible reagents and equipment. **(a)** Comparison of Genscript HDRTs with internally generated HDRTs for both ssCTS (top) and dsCTS templates (bottom). Shown for each are %knock-in, live cell counts, and knock-in cell counts in comparison to internally generated ssDNA or dsDNA controls, respectively. **(b)** Flow plots for T cell immunophenotype analysis based on CD45RA and CD62L expression at Day 7 and Day 10 post-activation. **(c)** Gating strategy for extended flow cytometric analysis demonstrating CD45RA+CD62L+CD45RO-CCR7+CD95+ population immunophenotypically consistent with a Tscm population. **(d)** Live cell counts for large-scale GMP-compatible manufacturing process at Day 7 and Day 10 post-activation. Each experiment was performed with T cells from 2 independent healthy human blood donors. Error bars indicate standard deviation. RNP = Ribonucleoprotein, CTS = Cas9 Target Site, dsCTS = dsDNA HDRT + CTS sites, ssCTS = ssDNA HDRT + CTS sites, HDRT = homology-directed-repair template, M = M3814, MT = M3814 + TSA, MTX = M3814 + TSA + XL413.

## Acknowledgments

We thank all members of the Marson laboratory for their thoughtful input and technical assistance. We thank Stacie Dodgson, Sarah Pyle, Fyodor Urnov, Bruce Schaar, Jon Woo, Jennifer Okano, and Jackie Sawin for their helpful suggestions and generous assistance. This research was supported by the NIAID (P01AI138962), the Multiple Myeloma Translational Initiative (MMTI) at UCSF, and the Innovative Genomics Institute. B.R.S was supported by the UCSF Herbert Perkins Cellular Therapy and Transfusion Medicine Fellowship, the CIRM Alpha Stem Cell Clinic Fellowship, and an NIH LRP grant from the NCATS. D.N.N. is supported by NIH grants L40AI140341 and K08AI153767 and the CIRM Alpha Stem Cell Clinic Fellowship. J.W. was supported by the Multiple Myeloma Translational Initiative (MMTI) at UCSF. F.B. was supported by the Care-for-Rare Foundation and the German Research Foundation (DFG). T.L.R. was supported by the UCSF Medical Scientist Training Program (T32GM007618), the UCSF Endocrinology Training Grant (T32 DK007418), and the NIDDK (F30DK120213). A.M. holds a Career Award for Medical Scientists from the Burroughs Wellcome Fund, is an investigator at the Chan Zuckerberg Biohub, and is a recipient of The Cancer Research Institute (CRI) Lloyd J. Old STAR grant. The Marson lab has received funds from the Innovative Genomics Institute (IGI), the Simons Foundation, and the Parker Institute for Cancer Immunotherapy (PICI). The Eyquem Lab has received funding from PICI and the Grand Multiple Myeloma Translational Initiative. M.R.M is funded by the Cancer Research Institute and the Human Vaccines Project Michelson Prize for Human Immunology. The UCSF Flow Cytometry Core was supported by NIH S10 RR028962 and the James B. Pendleton Charitable Trust. This research was made possible by a grant from the California Institute for Regenerative Medicine (Grant Number INFR-10361).

## Author Contributions

B.R.S, V.V., J.H.E, and A.M. designed the study. B.R.S, V.V., and A.H. performed ssCTS experiments. B.R.S and V.V. performed inhibitor experiments. B.R.S. and A.H. performed ORF replacement experiments. B.R.S and V.V. performed GMP-compatible manufacturing experiments. B.R.S., V.V., J.Y.C., A.T., J.E., J.H.E., T.G.M., and J.W. designed and performed BCMA-CAR experiments. D.N.N performed HSC experiments. Y.Y.C. and F.B. performed pooled knock-in experiments. S.V and M.R.M performed NK cell experiments. L.Y. designed and coordinated the large-scale production and downstream purification process of single-stranded DNA repairing template; H.L. supervised the regulatory requirements and QC methods for ssDNA. B.R.S., V.V., and A.M. wrote the manuscript with input from all authors.

## Data Availability Statement

Sequences for gRNAs and primers are provided in Supplementary Table 2. Plasmids containing the HDR template sequences used in the study will be made available through AddGene, and annotated DNA sequences for all constructs are available upon request. Flow cytometry raw data files are available upon request.

